# Novel Pipelines to Extract Differences in Proteome Dynamics Based on Health Status

**DOI:** 10.1101/2025.02.01.636015

**Authors:** Bowen Xu, Jiaying Zhao, Tianhui Huang, Sewanou Hermann Honfo, Caroline Trumpff, Martin Picard, Alan A. Cohen, Molei Liu

## Abstract

Understanding dynamics and co-regulatory patterns in the human proteome is a promising path for unraveling the molecular basis of health and disease. Nevertheless, there remains an open challenge in extracting concise information from high-throughput proteomic data that can effectively characterize and predict health. We develop novel statistical and computational pipelines to tackle this problem in a longitudinal saliva proteomics data set collected throughout the awakening response in six healthy controls and six subjects with severe mitochondrial disease (MitoD), a clinical condition caused by genetic mitochondrial defects that affects cellular energy transformation and alters multiple dimensions of health.

We undertook three independent unsupervised approaches to characterize proteome dynamics and assessed their ability to separate MitoD individuals from controls. First, we designed a permutation test to detect the global difference in the proteomic co-regulation structure between healthy and unhealthy subjects. Second, we performed non-linear embedding and cluster analysis on elasticity to capture a more complicated relationship between health and the proteome. Third, we developed a machine learning algorithm to extract low-dimensional representations of the proteome dynamic and use them to cluster subjects into healthy and unhealthy groups without any knowledge of their true status. All three methods showed clear differences between MitoD individuals and controls.

Our results revealed a significant and consistent association between MitoD status and the saliva proteome at multiple levels during the awakening response, including its dynamic change, co-regulation structure, and elasticity. This connection is not restricted to a few MitoD-specific proteins but spreads over a wide range of proteins from many body functions and pathways. Pipelines such as those shown here are the first step toward establishing interpretable and accurate prediction rules for health based on proteome dynamics.

## 1 Introduction

There is increasing recognition that human health depends on numerous interconnected physiological mechanisms that jointly regulate organismal responses to changing internal and external conditions [26, 7]. Although there are undoubtedly specific disease-causing mechanisms that can lead to specific pathologies, shared risk factors for many common chronic diseases and increasing evidence of interconnected physiological systems strongly suggest that a generalized phenomenon of physiological health, here termed ’intrinsic health’, could be conceptualized and quantified.

Currently, there is no broad framework to empirically and objectively quantify intrinsic health; achieving such a framework would have enormous benefits to our ability to quantify global health status, compare various interventions, and measure the health of populations. Previous authors [34, 1] have suggested that a stimulus-response paradigm and/or dynamic data are essential for measuring broad-spectrum health proxies such as frailty, resilience and robustness; arguably, the same should be true of intrinsic health. Specifically, if there is a quantifiable organism-level phenomenon of intrinsic health, we would make two predictions: (1) Response to a stressor (or, more broadly, changing conditions) should reflect not only a few molecules, mechanisms, or pathways, but should be reflected in broad changes across many systems; and (2) the overall dynamics of the system should differ systematically and markedly between healthy and unhealthy individuals.

Proteomics has emerged as a promising approach for characterizing such systemic responses, offering a comprehensive view of the molecular landscape underlying health and disease. Largescale studies [11] have demonstrated the power of proteomics to identify biomarkers and pathways for disease discrimination and therapeutic targets. By capturing the dynamic interplay of proteins across biological systems, proteomic data serves as an important resource to quantify and understand intrinsic health at the molecular level.

Here, we provide a preliminary test of these two hypotheses with an eye toward the eventual development of measures of intrinsic health. Specifically, we use a subsample of 12 individuals from the Mitochondrial Stress, Brain Imaging, and Epigenetics study (MiSBIE) – six individuals with severe mitochondrial disease caused by genetic lesions (MitoD) and six healthy controls [20]. Each individual was monitored across the awakening response, with saliva samples taken at four time points on two different days. The awakening response is a well-characterized circadian event and represents a strong candidate for observing a dynamic, multi-systemic response [31]. Each sample was analyzed for about 3000 proteins. We then constructed three alternative, unsupervised pipelines to examine how the proteome changes across the awakening response, and compared results to see if there were differences between healthy and unhealthy individuals.

## 2 Methods

### 2.1 Data description and preprocessing

The Mitochondrial Stress, Brain Imaging, and Epigenetics (MiSBIE) study is a study of 70 control subjects and 40 subjects with MitoD, designed to understand how energetics and mitochondrial function mediate relationships between the mind and body. Subjects were aged 18-60 (mean age = 38) and were 69% female. Two types of genetic mitochondrial lesions were present in the subjects: (1) the most common mtDNA point mutation, m.3243A>G, present in 25 subjects; and (2) large scale mtDNA deletions (15 subjects). All subjects underwent an extensive protocol that included baseline measurements, responses to various stressors, and multiple-day home-based assessment [20]. Here, we utilize proteomics data in home-collected saliva samples across the awakening response in a subsample of 12 individuals, 6 individuals with severe MitoD as the unhealthy group and 6 subjects without MitoD and matched with the disease cases by age and gender. The efficacy of saliva samples in reflecting the systemic physiological profile has been proven in earlier studies [30, 32, 17]. Following the Biomarker Network recommended procedure [21], saliva samples (1-2 *mL*) were collected at four time points t, including 0 minutes (denoted as t = 1), 30 minutes (t = 2), and 45 minutes (t = 3) post awakening, as well as before the bed time in the evening (t = 4, also denoted as pm). Each individual had saliva samples collected on three days; here, we analyzed samples from two of these days chosen randomly. We aim to use these response data to record changes in the proteome profile related to the awakening response, which is thought to involve comprehensive molecular biological processes in the human body [5]. An overview of the collection and the structure of our data is given in Fig. 1a.

**Figure 1:**
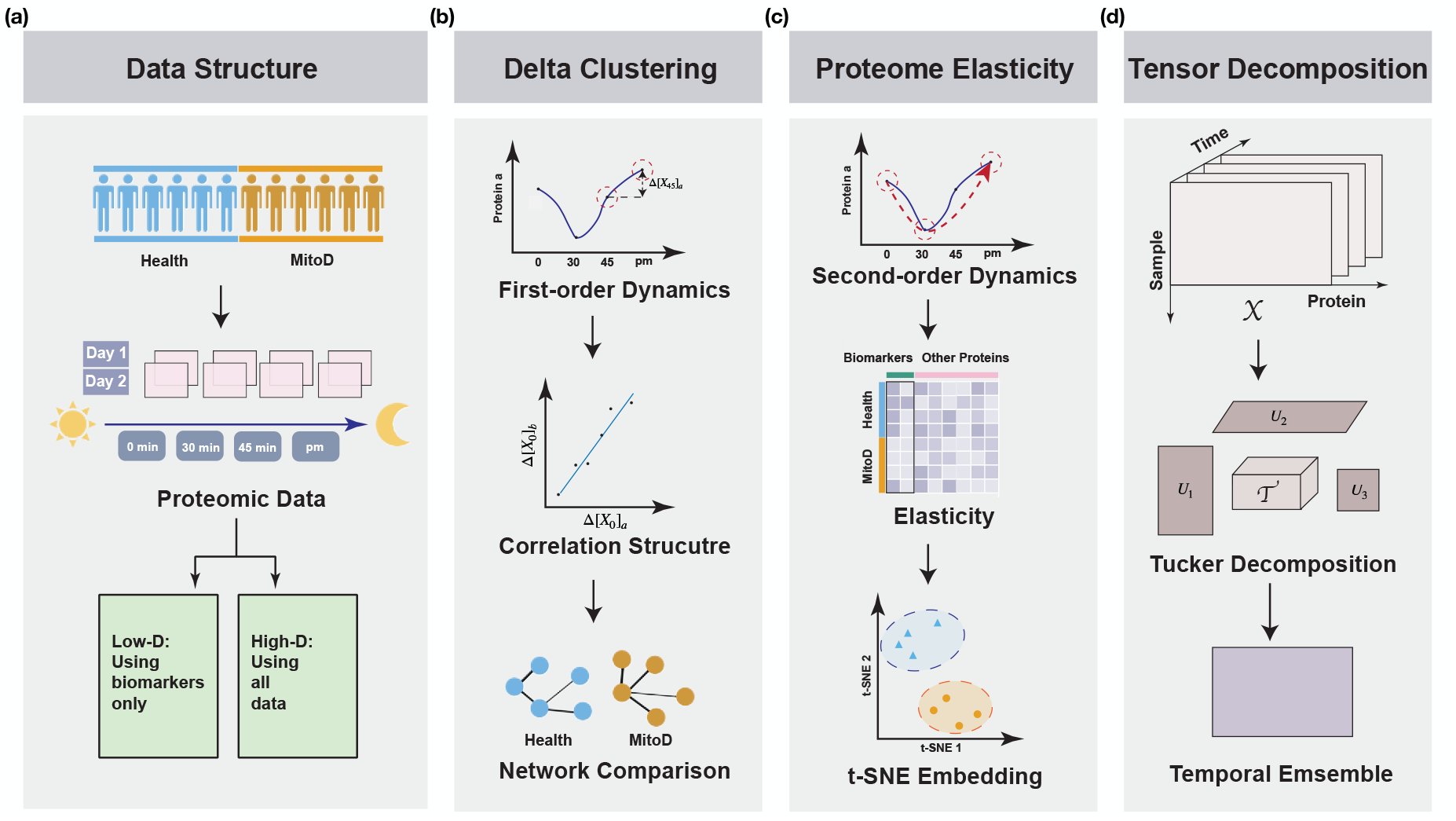
Overview of data structures and pipeline approaches. a. Data structure and broad approaches. b. Delta clustering pipeline (correlated changes). c. Proteome elasticity pipeline. d. Tensor decomposition pipeline.

Proteomics were quantified by Olink using their Explore 3072 platform, which utilizes proximity extension assays [25]. The protein expression level is quantified by the Normalized Protein expression (NPX) value using the technology developed by Olink. The NPX metric is derived from the cycle threshold (Ct) value and is presented on a log_2_ scale. Hereafter, we use **X** to denote sample-by-protein expressions, use superscript to denote the health (MitoD) status with *h* indicating healthy (i.e., not a MitoD case) and *uh* indicating unhealthy, and use subscript for time. For instance, 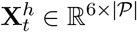 represents the proteomic profiling matrix for healthy samples at time *t*, where |𝒫| represents the number of proteins. We filtered out 207 proteins with either a high assay warning rate (>30%) or a high sample warning rate (50%) appearing due to technical reasons; the remaining |𝒫| was 2865 proteins.

### 2.2 Overview of goals and methods

Here, we introduce the general idea and outline of our statistical analyses. Our first goal was to test whether suites of correlated changes in the proteome during the awakening response differ by health (MitoD) status. To do this, we calculated the delta (change) of each protein across time points, and then assessed the correlation structure of these deltas - in other words, when one protein changes during the awakening response, which other proteins change with it? We performed this separately in healthy and unhealthy subjects, and then used cluster analysis to assess the extent to which these correlation structures differ. We compare the healthy-unhealthy separation with random separations to assess whether our differences exceed those expected due to chance. We call this approach “delta clustering.” Compared to simply testing whether protein concentrations differ between the groups (i.e., marginal mean differences), this approach should bring more insight into proteome dynamics (Fig. 1b).

Second, we go beyond the simple contrasts between two time points and consider the individuals’ elasticity to awakening reflected by their proteome. This is motivated by a recent study defining gene elasticity and profiling it across tissues in mice and monkeys [37]. Elasticity is determined by at least three time points before, during, and after a stimulus considered, the awakening response in our case. High elasticity characterizes the ability to both respond to and recover from a stimulus.

Considering both the nonlinearity and high-dimensionality of the proteome, we applied unsupervised manifold learning [33] to extract global patterns in elasticity across the entire proteome. We then projected the extracted features to assess whether variation in elasticity across the proteome in our sample was closely aligned with health status. We call this approach “proteome elasticity” (Fig. 1c).

Our final goal was to test whether there are broad differences by health status in the overall dynamic of the proteome across the four time points within a day. This required assessing variation across three types of variation - temporal, protein-level, and individual-level. We used tensor decomposition to separate these three types of variation and then focus on how the protein-level variation was structured within temporal variation that contrasted across timepoints. In other words, we wanted to extract and select low-dimensional representation for the proteome contrasting different time points in a purely data-driven way. We then assessed whether these time-varying proteomic structures were associated with health status (again, an unsupervised approach). We call this approach “dynamic tensor decomposition” (Fig. 1d).

To investigate and validate the informativeness of the proteomic analysis involving so many proteins not specific to MitoD, we identified *GDF15* and Interleukin 6 (*IL6*), two important biomarkers of MitoD found in earlier studies [24, 4]. We properly adapted all the analyses mentioned above to low-dimensional (low-D) scenarios only involving *GDF15* and *IL6*. This was to see if our developed pipelines were able to extract stronger and more generalizable signals for health, compared to lower dimensional and more disease-specific biomarkers. Our hypothesis was that the high-dimensional proteomic analyses could differentiate healthy and unhealthy groups more clearly and effectively than targeted, low-dimensional ones. In the following sections, we present the pipelines we developed and results for the three goals one after another.

### 2.3 Comparing co-regulation networks of expression dynamics in health and MitoD systems

In the first pipeline, our goal was to test whether suites of correlated changes in the proteome during the awakening response differ by health (MitoD) status. We first employ a finite difference method to characterize protein dynamics. To be specific, we use the first order difference (delta) of protein expressions (Δ[**X**_*t*_] := **X**_*t*+1_ − **X**_*t*_, *t* = 1, …, 3) along the time scale to approximate the protein velocity, i.e., the time derivative of protein concentration in the studied physiological system. For instance, Δ[**X**_*t*_]_*sl*_ measures the trend of protein *l*’s expression level in sample *s* from time *t* to *t* + 1. Then we explore the correlation structure of dynamic variants by constructing co-regulation networks 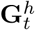 and 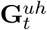 from those computed deltas for health and MitoD systems at time *t* (*t* = 1, …, 3). Let 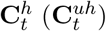 be the corresponding adjacency matrix for 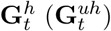. At time *t*, the co-regulation relationship for health and MitoD samples is quantified by Pearson’s coefficient, denoted by 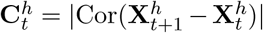 and 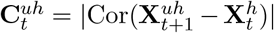, where Cor(·) denotes correlation matrix and | · | represents absolute value. Moreover, close to 1 indicates that the expression trends of protein *i* demonstrate strong correlation, either positive or negative, with the trends of protein *j* during the awakening response in a healthy system, while 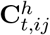 close to 0 suggests the dynamics of protein *i* and *j* have weak associations. Compared to simply testing whether protein concentration deltas differ between the groups (i.e., marginal mean differences), we aimed to assess the global difference of co-regulation networks 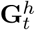 and 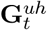, which should bring more insight into proteome dynamics. Therefore, we conduct the following hypothesis test,

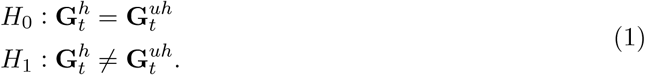

To do this, we define a distance measure to quantify the discrepancy between 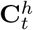 and 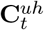. Due to the noise arising from the high-dimensionality, we first trim the difference between 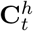 and 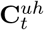 by removing entries with absolute values smaller than some threshold parameter *λ*. Let TrimDiff(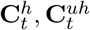; *λ*) be the trimmed matrix with its (*i, j*)-th entry computed by

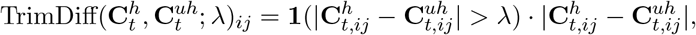

where **1**(·) represents the indicator function. Then we use its Frobenius norm for testing and apply a permutation approach [16] to determine statistical significance of co-regulation discrepancy

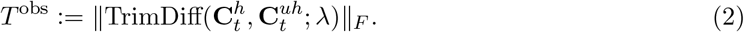

Under the null hypothesis, protein distributions of healthy and MitoD populations are identical so that labels of the two categories are exchangeable. Hence, we design the permutation method in the following way. We randomly assign the MitoD label to half of the individuals denoted as *r*_1_, as well as the healthy label to the other half *r*_0_, and compute the network discrepancy for permutated samples as

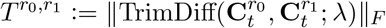

where 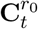 and 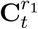 are obtained in a similar way to the observed samples. When there is an essentially larger difference between the healthy and unhealthy groups, the observed *T* ^*obs*^ is expected to be significantly larger than 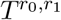. We obtain the p-value by the following procedures,

**S.1** For the 12 individuals, divide them into *r*_0_ and *r*_1_ of an equal size 6.

**S.2** Calculate 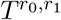.

**S.3** Conduct steps 1 and 2 for all 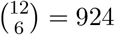 combinations.

**S.4** Compute the p-value as the proportion of assignments with 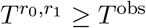.

### 2.4 Higher-order characterization of dynamics through proteomic elasticity

In our second pipeline, we go beyond the first-order expression difference between two consecutive time points and consider a second-order method in characterizing proteome dynamics, “protein elasticity”. Motivated by a recent study defining gene elasticity across tissues in mice and monkeys [37], we formulate protein elasticity to profile three time points before (*t* = 0), during (*t* = 30), and after a stimulus considered (*t* = *pm*, see Fig. 1c for illustration).

The elasticity of a protein *j* for sample *s* in response to awakening is computed as:

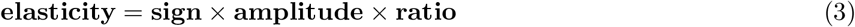

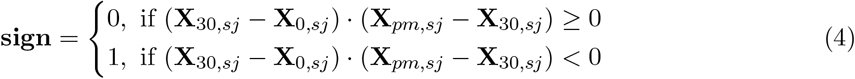

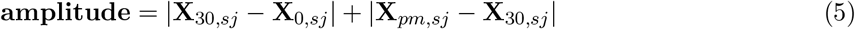

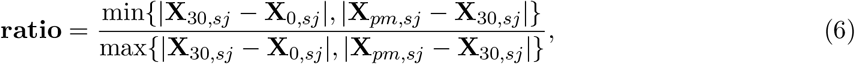

where we aim to measure the capacity of the human body to cope with and recover from external stimulus. There are three key components in this definition including the **sign, amplitude** and **ratio**. The **sign** ensures that the computed elasticity is non-zero only when there is a recovery process such that the expressions across 0 min (*t* = 1), 30 min (*t* = 2), and pm (*t* = 4, bedtime) form a U-shaped or inverse-U-shaped trend (see Fig. 1c for instance) rather than a linear trend, i.e., successive increase or decrease. To describe the biomarker’s deviation caused by perturbations, in our case, awakening, we compute the total variations as **amplitude** to measure the magnitude of oscillations. Defined by the fraction of relative changes between |**X**_30,*sj*_ −**X**_0,*sj*_| and |**X**_*pm,sj*_ −**X**_30,*sj*_|, the **ratio** lies within [0, 1] and quantifies the resilience of the studied physiological system. A high **ratio** indicates a protein’s ability to both respond to and recover from a stimulus. For each sample *s* ∈ {1, …, |𝒮|} and protein *j* ∈ {1, …, |𝒫|}, we use the product of **sign, amplitude**, and **ratio** to characterize the overall elasticity, and obtain a proteomic elasticity matrix denoted as **E** ∈ ℝ^|𝒮|×|𝒫|^, where **E**_*sj*_ is computed by Eq. (3). We first use the biomarker elasticities to test whether the low-dimensional data is informative enough to reveal the systemic-level health condition. To select appropriate biomarkers, we retain proteins with high mean and variance in elasticity results. Then we propose to use t-distributed Stochastic Neighbor Embedding (t-SNE) [33] on **E** to investigate whether protein elasticity provides more insights in distinguishing health status. t-SNE is a nonlinear embedding method that quantifies elasticity similarities for each sample pairs through the kernel of the t-distribution. By implementing t-SNE, we project samples onto a two-dimensional latent space based on computed elasticities. The proximity of data points in the embedded space reflects the degree of similarity in their elasticity profiles.

### 2.5 Temporal representation learning based on tensor decomposition

To assess expression dynamics over three directions, i.e. time-level, protein-level, and individual-level variation, we reformulated our data into a 3rd-order tensor by concatenating sample-by-protein matrices along the time mode. Let **𝒳** ∈ ℝ^|𝒮|×|𝒫|×4^ be the tensor that denotes the stacked expression matrices (see Fig. 1d for illustration), where |𝒮| represents sample number and |𝒫| represents the number of filtered proteins. We use a colon to indicate all elements of a mode such that the slice **𝒳**_::0_ := **X**_0_ represents protein expressions in all samples at *t* = 0. Our primary aim is to condense high-dimensional and dynamic proteomic expression into a few representative features that are predictive of health. Rather than employing traditional representation learning methods like principal component analysis (PCA), which fails to tackle higher-order tensors, we propose to implement Tucker decomposition [22] to address this 3-dimensional array **𝒳**. A toy model illustration is given in Fig. 1d. Denote **𝒯** ∈ ℝ^3×10×3^ as the core tensor. Then we can decompose **𝒳** as Eq.(7)

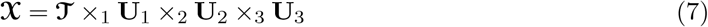

where ×_*k*_, *k* = 1, …, 3 indicates *k*-mode product and **U**_1_ ∈ ℝ^|𝒮|×3^, **U**_2_ ∈ ℝ^|𝒫|×10^ and **U**_3_ ∈ ℝ^4×3^ are factor matrices for each mode satisfying

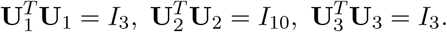

Specifically, the core tensor **𝒯** can be regarded as a compressed version of **𝒳**. Similar to PCA, the factor matrix can be treated as a linear operator that maps **𝒳** onto a new coordinate system that can capture representative features of **𝒳**. Given factor matrix **U**_3_, we obtain higher-order principal components for tensor **𝒳** in the time-mode as 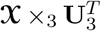. Each 2-order principal component can be regarded as a temporal ensemble of **X**_*t*_ (*t* = 1, …, 4) by using column vectors of **U**_3_ as weights. Suppose *w*_1_, …, *w*_4_ are one set of weights (a row of **U**_3_); we then compute temporally aggregated matrix 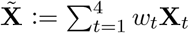 to obtain an average representative of protein expressions along the time mode. Here, 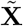 can be used to encapsulate the dynamic equilibrium characteristics inherent in the proteome. Moreover, as a higher-order principal component for **𝒳**, the matrix profile 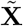 may aid in health status inference. Hence, we further perform PCA to extract the top 10 principal components (PCs) of the projected proteome profiles 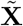, and denote the embedded ten features as **V** ∈ ℝ^|𝒮 |×10^. We then visualize the low-dimensional representation (columns of **V**) and compare them with the health (MitoD) status. We then select three PCs among the ten that are most capable of separating subjects into two groups by conducting unsupervised discriminative feature selection [36] (UDFS). UDFS is a linear discriminator that uses a convex regularizer (penalty) for variable selection, which aims to select a subset of features for compact and accurate data representation. We also consider applying non-convex UDFS [38], a generalized UDFS method that utilizes non-convex penalty, for example *L*_2,*p*_ (0 *< p <* 1) norm, for feature selection. Finally, we use a 3D scatter plot to present the three selected PCs to assess whether the aggregated profile 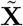 improves our understanding of health status.

## 3 Results

### 3.1 Systems-level difference of health and MitoD status during the awakening response

In pipeline 1, we focused on characterizing the difference between health and MitoD status during the awakening response at the systems-level from a network perspective. As Fig. 1b illustrates, we first created co-regulation networks 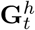 and 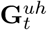 that reflect dynamic protein associations for health and MitoD systems respectively based on Pearson’s correlation of the first-order expression difference Δ[**X**_*t*_]. Then we (i) conducted hypothesis testing (1) to assess the global difference between the two networks through the trimmed matrix discrepancy norm Eq.(2); and (ii) evaluated whether low-dimensional biomarkers are informative enough to reveal the health status of the physiological system.

Since we assume that proteins with low variation across the time course are non-informative, we first selected the 2500 most dynamic proteins with the largest coefficient of variation (CV) on deltas and obtain a filtered expression matrix **X**_*t*_ ∈ ℝ^|𝒮 |×2500^.

We focus on the time interval around *t* = 1 to investigate regulation differences from just waking up to 30 minutes post-awakening, and conduct permutation tests on the global proteome network (2500 nodes) and biomarkers (sub-network between two biomarkers and the other 2500 proteins) respectively by **S.1**-**S.4**. For sensitivity analyses, we implemented various *λ* including 0.1, 0.2, and 0.3 to evaluate whether permutation results are affected by the choice of threshold *λ*. The p-values for the proteome network under *λ* = 0.1, 0.2, 0.3 are all 0.0411 (*<* 0.05), indicating that proteome co-regulation networks demonstrate significant differences in the two health states and that the results are insensitive to the choice of threshold *λ*. P-values are identical because the number of permutations is limited and an identical number of permutations were more extreme than the observed *T* ^*obs*^. In contrast, the p-values for low-dimensional (i.e., *GDF15* and *IL6*) subnetwork tests are all 1 (> 0.05) for *λ* = 0.1, 0.2, 0.3. Therefore, using the whole proteome network appears to improve our understanding of intrinsic health beyond simply using previously identified biomarkers.

To give an intuitive illustration of the permutation approach, we use heatmaps of the (permutated) proteome co-regulation networks at *t* = 1 (Fig. 2). After filtering proteins with low expression mean and variance, we retain 421 highly variable proteins for the heatmap example. Let 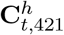 be the curated network for healthy samples and 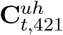 be its corresponding co-regulation network in unhealthy samples. After performing permutation to re-assign health status labels, we use 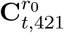 and 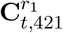 respectively to denote co-regulation networks for the permutation results. Rows and columns of 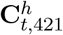 and 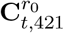 are re-arranged by hierarchical clustering (number of clusters *n* = 5). For 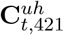 and 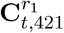, the orders of their rows (columns) are in accordance with 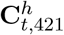 and 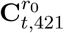, respectively. Note that the cluster structure of 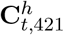 (Fig. 2a) appears completely absent in 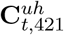 (Fig. 2b), whereas the cluster structure of 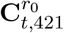 (Fig. 2c) is at least partially replicated in 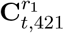 (Fig. 2d), despite having been derived from only six individuals.

**Figure 2:**
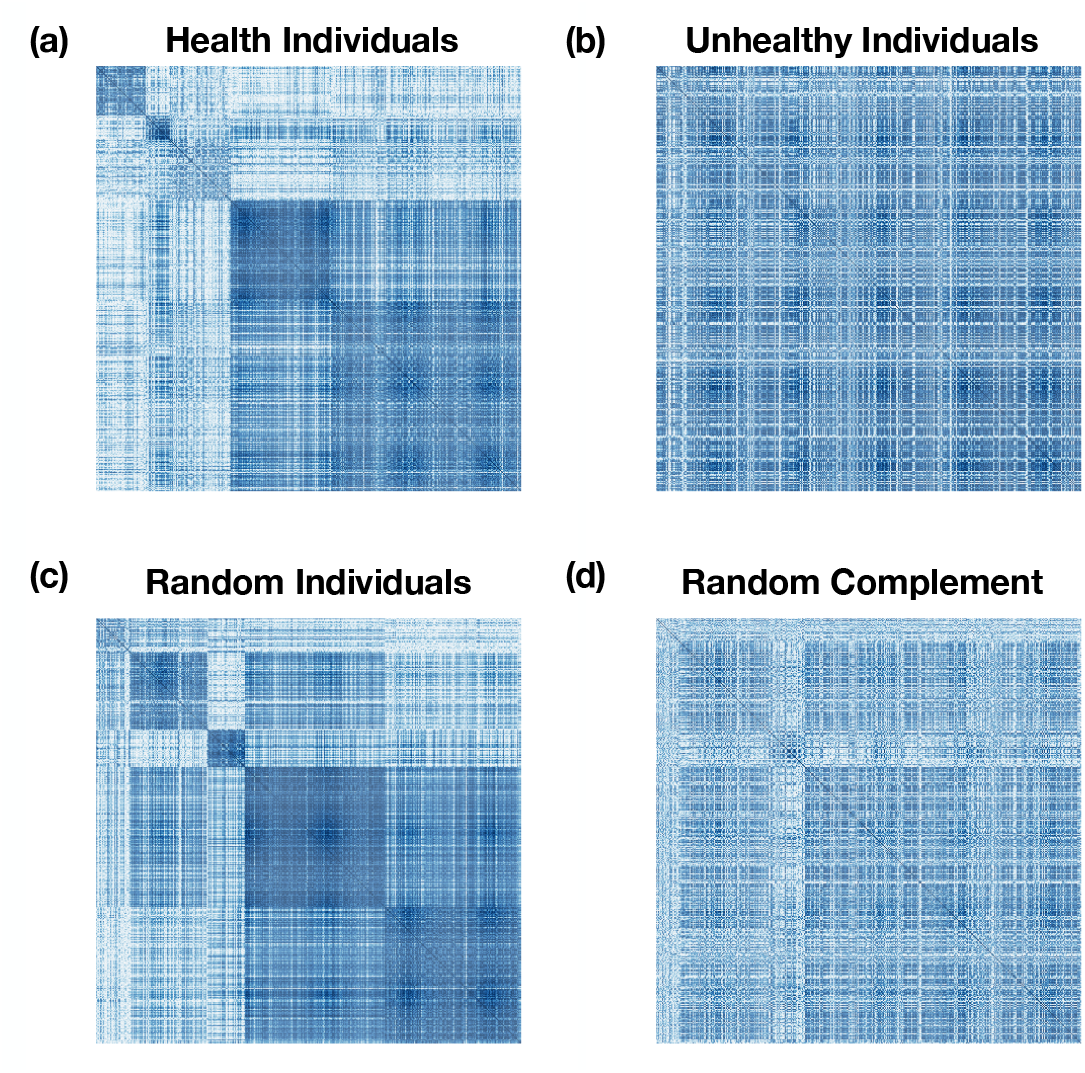
(a) Heatmap of 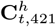. (b) Heatmap of 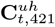. (c) Heatmap of 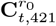. (b) Heatmap of 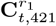.

### 3.2 Nonlinear embedding of proteomic elasticity

As illustrated in Section 2.4, we first obtained elasticity matrix **E** and focused on the elasticity profiles of biomarkers *IL6* and *GDF15*. We provide the bar plot (Fig. 3a) of mean elasticity in Healthy and MitoD samples for *IL6* and *GDF15* respectively, where the error bar represents standard error of the mean (SEM). Mean elasticity differs between the two health conditions, especially in the case of *GDF15* (Healthy 0.114596, MitoD: 0.608519), which implies that *GDF15* may be markedly different during the awakening process in diseased physiological systems. However, the relative mean values of biomarker elasticity do not consistently align with health states; for instance, the mean elasticity of *IL6* is higher in healthy samples compared to unhealthy ones, whereas the opposite is observed for *GDF15*. Accordingly, we examined other specific proteins chosen via data-driven methods to see if low-dimensional biomarker elasticity was capable of predicting health status. To do this, we retained proteins for which no more than four individuals had zero elasticity (137 proteins excluded). Among these, we chose the two proteins with the largest variance, *LSM1* and *MEGF9*. A scatter plot was then generated to visualize the elasticity results for these biomarkers. As illustrated in Fig. 3b, healthy samples consistently exhibited low elasticity in both *LSM1* and *MEGF9*, while MitoD samples displayed a wide range of elasticity values. Specifically, half of the diseased samples had very low elasticities, overlapping with healthy samples, while the other half demonstrated significantly higher elasticity values. While this pattern suggests that healthy and unhealthy individuals may differ in their elasticity profiles on *LSM1* and *MEGF9*, it also implies that the elasticities of low-dimensional biomarkers will not be able to clearly distinguish healthy from unhealthy (Fig. 3b).

**Figure 3:**
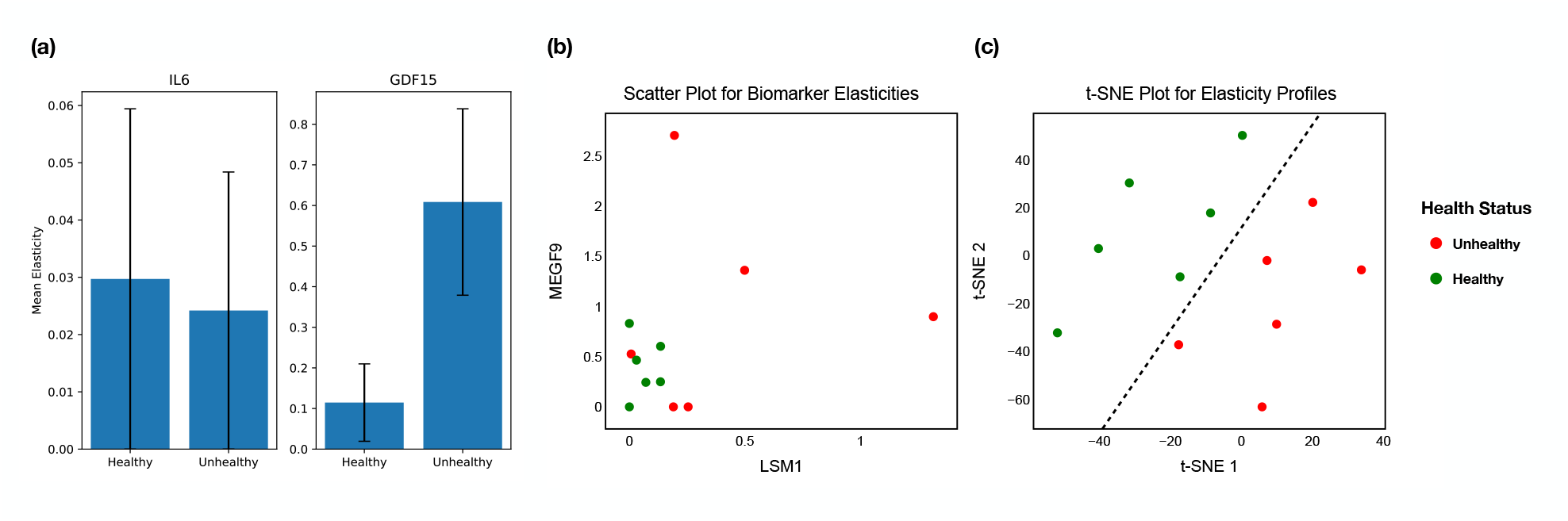
Elasticity: (a) Barplot of biomarker elasticities; error bars indicate standard error of the mean (SEM). (b) Scatter plot of selected biomarker elasticities. (c) t-SNE plot for **E**.

To overcome the limitations of using a small number of pre-identified biomarker profiles, we therefore studied the transformed data (elasticity) from a systems’ perspective to assess whether the joint utilization of the entire proteome improves our ability to distinguish healthy from unhealthy. We performed t-SNE on **E**, the global elasticity profile, and provide the results in Fig. 3c, where we set 10 as the number of nearest neighbors, choose 3 for the number of PCs and set 1500 as the number of maximum iterations. Fig. 3c gives a well-separated rather than overlapping low-dimensional representation and demonstrates that we can classify healthy and diseased samples through a separating hyperplane (see the dashed line), which suggests that the global elasticity profile contains substantial information reflecting population health status. We also conduct sensitivity analysis on the number of iterations (*n* = 500, 600, 800, 1500, 3000, 4000, 5000) to investigate whether optimization affects visualization results; the corresponding t-SNE plots are given in Supplementary File Fig. S1.

### 3.3 Dynamic tensor decomposition: Representation learning and clustering on proteomic dynamics

In this section, we start by implementing Tucker decomposition on **𝒳** and present the result of **U**_3_, the temporal dimension, in Table 1. The first factor of **U**_3_ indicates a global up-regulation or down-regulation across time, while the second factor demonstrates contrasts to baseline (awakening) from subsequent timepoints, and the third factor is harder to interpret. The first factor is not of interest in this context, as we wish to examine dynamic patterns, i.e., those that vary temporally. Accordingly, we focus our attention on the second factor, which has a clear biological interpretation. We then apply *U*_3_ on **𝒳** to obtain higher-order principal components 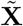 along the second factor of the time mode. Let ***u***_31_, ***u***_32_ and ***u***_33_ denote columns of **U**_3_, then we can obtain 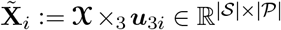 as 2nd-order principal components. We first study whether variation in the pre-selected biomarkers, *GDF15* and *IL6*, along 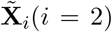, the linearly transformed data of **𝒳**, is informative enough to reflect health status. In Fig. 4a, we provide the scatter plot of the averaged expressions of *IL6* and *GDF15* for each sample in 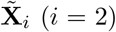. It is evident that there is no clear separation between healthy and MitoD individuals.

**Table 1:** Temporal weights fitted by tensor decomposition.

|  | 1st factor | 2nd factor | 3rd factor |
| --- | --- | --- | --- |
| 0 min | 0.83 | -0.55 | -0.09 |
| 30 min | 0.32 | 0.56 | -0.63 |
| 45 min | 0.36 | 0.41 | 0.77 |
| pm | 0.27 | 0.47 | -0.02 |

**Figure 4:**
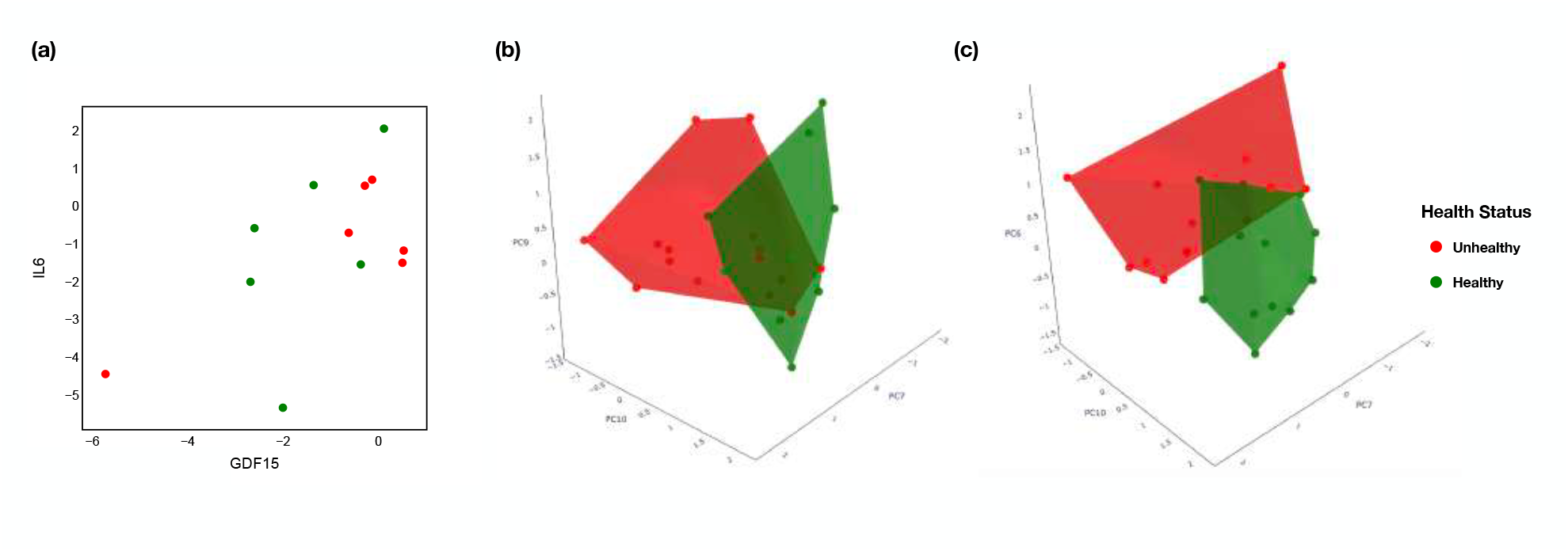
Tensor analysis: (a) Temporal patterns of *GDF15* and *IL6* along the second tensor fail to distinguish healthy from MitoD individuals. (b) 3D visualization of 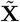 along (***v***_7_, ***v***_9_, ***v***_10_) showing reasonable but not perfect separation of healthy and MitoD individuals. (c) 3D visualization of 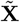 along (***v***_6_, ***v***_7_, ***v***_10_) showing reasonable but not perfect separation of healthy and MitoD individuals.

Next, we incorporate variation in the full proteome along 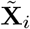 to investigate whether the high-dimensional profile of 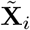, the linear projection of **X**_*t*_ (*t* = 1, …, 4) along the time direction, is able to reveal the intrinsic health state. As described in Section 2.5, we perform PCA on 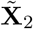 and employ UDFS and non-convex UDFS to select informative PCs. Let ***v***_*j*_ (*j* = 1, …, 10) denote the *j*-th PC. The two UDFS approaches select (***v***_7_, ***v***_9_, ***v***_10_) and (***v***_6_, ***v***_7_, ***v***_10_) respectively, with two PCs, ***v***_7_ and ***v***_10_, selected by both methods. In Supplementary File Table S1, we calculate the proportion of the variation in 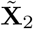 explained by the selected PCs and list the top 20 proteins (in their gene names) with the largest contributions on them. The top contributing proteins for each selected PC are passed over to a pathway enrichment analysis via shinyGO [15], with the detected pathways presented in Supplement File Table S2. The 3D plot of two sets of PCs are presented in Fig. 4b and c, with the healthy samples labeled in green and unhealthy individuals labeled in red. One can find a clear hyperplane separating the two groups, and the small overlap between the convex hulls of two populations is acceptable. As we can see from Fig. 4, using the whole proteome of 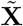 instead of two biomarkers chosen based on *a priori* knowledge improves the result of low-dimensional representations in terms of characterizing health.

## 4 Discussion

Here, we used three separate, unsupervised pipelines to characterize dynamics of the saliva proteome across the awakening response and compare them in healthy versus unhealthy individuals, defined by MitoD status. All three methods showed the ability to clearly distinguish by health status, despite the small sample size (six individuals each in healthy and unhealthy groups); all three clearly outperform what can be achieved with only targeted, mechanistically identified proteins. Broadly, these results strongly confirm both of our hypotheses: (1) that signals of the dynamics are not confined to one or a few mechanistically identifiable proteins, but are broadly shared across the circulating proteome, with stronger signals identified via broad approaches; and (2) that dynamics of the proteome across the awakening response are fundamentally different in individuals with and without MitoD – not only is it possible to distinguish health status (as might easily be shown via a supervised approach); differences in health status are so important that they emerge on their own as major drivers of variation in proteome dynamics, regardless of which of these three methods is used.

Our analysis here is intended to be a proof-of-principle rather than a definitive demonstration of the best possible pipelines. These particular pipelines may or may not be retained as optimal or effective once validated in a larger sample. Given the complexity of the data structure we were working with, many more pipelines and analytical approaches might be envisioned, and we strongly encourage other groups to propose other approaches using these data (available upon reasonable request from qualified researchers) or others with similarly appropriate structures. Nonetheless, the three approaches proposed here are conceptually distinct. The fact that all three work well with such a small sample is confirmation of the broad potential of such methods.

Numerous previous studies have examined how various -omics profiles respond to stressors. However, most of these studies are in model organisms such as yeast or bacteria [19, 3], or under environmental stress, such as crops or wildlife [23]. Relatively few have examined the dynamics of a response in real time. Such dynamics responses have been studied in drug-treated individual cancer cells [9], in plant responses to heat stress [12], and in growth and stess response of cell [28]. We are not aware of other studies examining the dynamics blood, urine, or saliva proteomics in response to acute stressors in humans. Accordingly, the methods for treating such data are also a work in progress. The pipelines proposed here may represent an important first step.

While the best way to characterize the response of an entire high-dimensional array of -omics data remains an unresolved problem, more progress has been made in measuring the response of single dynamic parameters. For example, the oral glucose tolerance test (OGTT) has long been used as a marker of metabolic health [18]. However, even in this case, the best way to model a dynamic response from a limited number of time points remains a challenging problem [29].

Here we have used saliva proteomic data. This is not the only data modality that may be relevant – metabolomics is also highly promising, for example [2]. Proteomics was chosen because it is likely to vary at a relevant timescale (as indeed was observed here) and is perhaps most directly related to the functional interpretation of the biology. Transcriptomics patterns, for example, do not always parallel proteomics patterns, and can be less stable across populations [27].

Future validation of these and similar pipelines will be essential, both in an expanded sampling of the cohort here and in other datasets stratified by other health conditions. Such health conditions could include, for example, frailty, diabetes, age-related decreases in oxidative phosphorylation capacity, hemodialysis, surgery recovery, and bone marrow transplant [14, 10, 13, 6, 35, 20]. Here, we have used the awakening response as a stimulus paradigm, but we predict that similar findings would emerge from any major stimulus-response paradigm operating at the organism level [34]. We hypothesize that, while each type of impairment via a specific health condition will have unique features, all impairments may also share common features that are evident in deviations from a healthy norm, a characteristic response of healthy individuals to a given stimulus. Future research should evaluate the extent to which such deviations are unique to specific health impairments, as well as the universality of a healthy norm across ages, sexes, and countries. In addition, it is interesting to further apply our pipelines to investigate other types of proteomic samples (e.g., those collected from blood) as well as other types of -omics data.

The results presented are, in this sense, a small first step towards a general characterization of what the normal proteome dynamics are in healthy individuals and how this changes as health is lost. Such a characterization will open the doors toward measurement of intrinsic health, and will thereby serve as a crucial tool to build a new science of health focused on prevention and maintenance of basic homeostatic states. The noninvasive and widely applicable nature of the saliva sampling method is also helpful for achieving this goal in practice. This would allow for a radical departure from our current, disease-focused paradigm, allowing treatments to emerge well before clinical signs of dysfunction appear as early deviations from such a healthy norm are detected [8].

