## Supplementary material for "Novel Pipelines to Extract Differences in Proteome Dynamics Based on Health Status"

February 1, 2025

### 1 Supplementary Tables

| Selected PCs | Explained Variation | Top 20 Contributive Proteins |
|---|---|---|
| PC 6 | 3.605429% | KIR3DL1, CDSN, SERPINB5, IGSF3, SPART, ANXA2, TNFSF10, CEP20, CKAP4, MICB_MICA, FCN1, SNX15, LCN15, GSTT2B, IL18, LCAT, TOMM20, MAPKAPK2, LGALS7_LGALS7B, LGMN |
| PC 7 | 2.778682% | NOS2, MMP10, LCN15, CERT, BAX, IL10, MICB_MICA, BTN1A1, SFTPA2, CTSF, LTBP3, POF1B, ZCCHC8, SCIN, NDUFS6, KLK14, LZTFL1, CD2AP, PSCA, CD63 |
| PC 9 | 1.958239% | PRTFDC1, BTN1A1, KLK12, NDUFS6, VTA1, MMP8, FIS1, FUT3_FUT5, PRDX6, SKAP2, MINK1, LGALS7_LGALS7B, LILRB2, CD207, BAIAP2, MYH9, KLK10, PSME1, NPM1, PACS2 |
| PC 10 | 1.713381% | NOS2, MMP10, LCN15, CERT, BAX, IL10, MICB_MICA, BTN1A1, SFTPA2, CTSF, LTBP3, POF1B, ZCCHC8, SCIN, NDUFS6, KLK14, LZTFL1, CD2AP, PSCA, CD63 |

Table S1: Results for Selected PCs by UDFS, Explained Variation and Top 20 Contributive Proteins

### 2 Supplementary Figures

| PC 6 | | | | |
|---|---|---|---|---|
| **Enrichment FDR** | **nGenes** | **Pathway Genes** | **Fold Enrichment** | **Pathways** |
| $1.8 \times 10^{-2}$ | 2 | 102 | 56.1 | Amoebiasis |
| $1.8 \times 10^{-2}$ | 2 | 151 | 37.9 | Phagosome |
| $1.8 \times 10^{-2}$ | 2 | 179 | 32 | Tuberculosis |
| $1.8 \times 10^{-2}$ | 2 | 249 | 23 | Salmonella infection |
| **PC 8** | | | | |
| **Enrichment FDR** | **nGenes** | **Pathway Genes** | **Fold Enrichment** | **Pathways** |
| $3.9 \times 10^{-2}$ | 2 | 137 | 33.4 | Yersinia infection |
| **PC 9** | | | | |
| **Enrichment FDR** | **nGenes** | **Pathway Genes** | **Fold Enrichment** | **Pathways** |
| $1.5 \times 10^{-2}$ | 2 | 72 | 79.4 | Mitophagy-animal |
| **PC 10** | | | | |
| **Enrichment FDR** | **nGenes** | **Pathway Genes** | **Fold Enrichment** | **Pathways** |
| $3.03 \times 10^{-3}$ | 2 | 37 | 154.6 | African trypanosomiasis |
| $3.03 \times 10^{-3}$ | 2 | 42 | 136.2 | Graft-versus-host disease |
| $3.03 \times 10^{-3}$ | 2 | 49 | 116.7 | Malaria |
| $4.7 \times 10^{-2}$ | 1 | 30 | 95.3 | Antifolate resistance |
| $4.0 \times 10^{-3}$ | 2 | 65 | 88 | Inflammatory bowel disease |
| $4.4 \times 10^{-3}$ | 2 | 92 | 62.2 | Rheumatoid arthritis |
| $4.4 \times 10^{-3}$ | 2 | 93 | 61.5 | IL-17 signaling pathway |
| $4.4 \times 10^{-3}$ | 2 | 101 | 56.6 | Chagas disease |
| $4.4 \times 10^{-3}$ | 2 | 102 | 56.1 | Amoebiasis |
| $4.4 \times 10^{-3}$ | 2 | 108 | 53 | Th17 cell differentiation |
| $4.4 \times 10^{-3}$ | 2 | 109 | 52.5 | HIF-1 signaling pathway |
| $8.4 \times 10^{-3}$ | 2 | 162 | 35.3 | JAK-STAT signaling pathway |
| $8.4 \times 10^{-3}$ | 2 | 169 | 33.8 | Protein processing in endoplasmic reticulum |
| $8.4 \times 10^{-3}$ | 2 | 171 | 33.5 | Influenza A |
| $8.5 \times 10^{-3}$ | 2 | 179 | 32 | Tuberculosis |
| $2.1 \times 10^{-2}$ | 2 | 294 | 19.5 | Cytokine-cytokine receptor Interaction |

Table S2: Enrichment Analysis Results for Selected PCs

| Elastic Biomarkers |
|---|
| LSM1, MEGF9, LMOD2, PODXL, PLB1, PFKFB2, GNLY, MGLL, CEACAM8, RET, ANXA2, MELTF, IFNL2, DTX3, TNFRSF17, CLMP, ACE, CD55, SERPINE2, CEACAM5 |

Table S3: Elastic Biomarkers

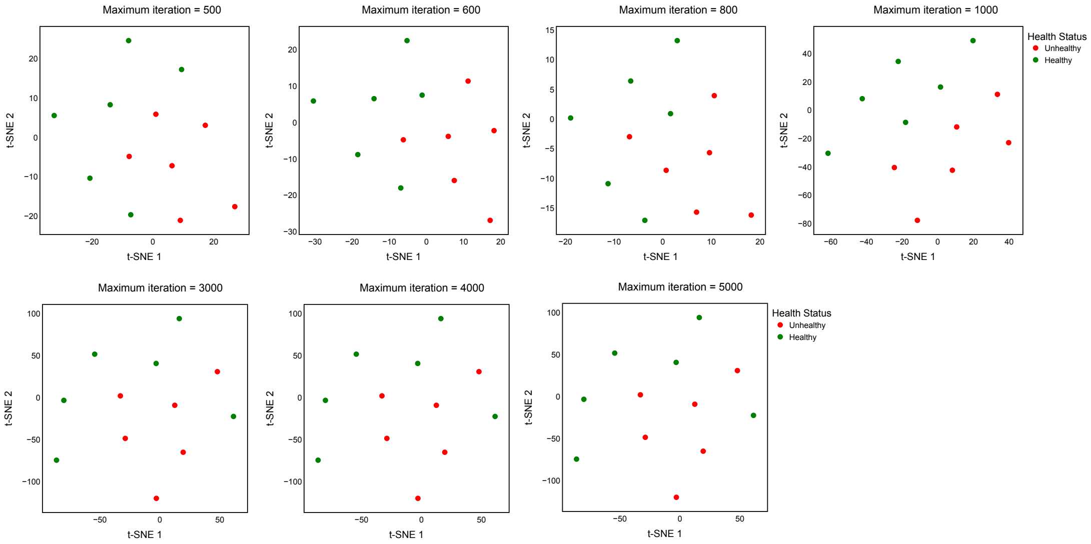


Figure S1: Sensitivity Analysis for t-SNE
